# Identification of NPC1 as a novel SARS-CoV-2 intracellular target

**DOI:** 10.1101/2020.12.19.423584

**Authors:** Isabel Garcia-Dorival, Miguel Ángel Cuesta-Geijo, Lucía Barrado-Gil, Inmaculada Galindo, Jesús Urquiza, Ana del Puerto, Carmen Gil, Nuria Campillo, Ana Martínez, Covadonga Alonso

**Affiliations:** Dpt. Biotechnology, Instituto Nacional de Investigación y Tecnología Agraria y Alimentaria (INIA), Ctra. de la Coruña km 7.5, 28040 Madrid, Spain; Centro de Investigaciones Biológicas Margarita Salas (CSIC), Ramiro de Maeztu 9, 28040 Madrid, Spain

**Keywords:** SARS-CoV-2, N protein, NPC1, target, antivirals

## Abstract

Niemann-Pick type C1 (NPC1) receptor is an endosomal membrane protein that regulates intracellular cholesterol trafficking, which is crucial in the Ebola virus (EBOV) cycle. The severe acute respiratory syndrome coronavirus 2 (SARS-CoV-2) enters the cell by binding of the viral spike (S) protein to the ACE2 receptor. This requires S-protein processing either by the surface transmembrane serine protease TMPRSS2 for plasma membrane fusion or cathepsin L for endosomal entry. Additional host factors are required for viral fusion at endosomes. Here, we report a novel interaction of the SARS-CoV-2 nucleoprotein (N) with the cholesterol transporter NPC1. Moreover, small molecules interfering with NPC1 that inhibit EBOV entry, also inhibited human coronavirus. Our findings suggest an important role for NPC1 in SARS-CoV-2 infection, a common strategy shared with EBOV, and a potential therapeutic target to fight against COVID-19.

## Introduction

To date, the COVID-19 pandemic has caused over one million of deaths and affected over 73-millions of people around the world (*1*). COVID-19 is caused by the emerging and pathogenic severe acute respiratory syndrome coronavirus 2 (SARS-CoV-2) first reported in the city of Wuhan (China) as a rare pneumonia (*2, 3*) that rapidly spread worldwide. A better understanding of the biology of SARS-CoV-2 is vital in order to develop effective therapeutics.

A key step of the biology of SARS-CoV-2 is the cell entry mechanism. Viral entry involves the activation of its trimeric spike glycoprotein by TMPRSS2 protease upon interaction with the angiotensin-converting enzyme 2 (ACE2) receptor at the plasma membrane to mediate fusion (*4-8*). Also, the virus can enter via endosomes under cathepsin L processing, similar to SARS-CoV-1 (*7, 9*). In fact, the inhibition of both alternative entries is necessary for full inhibition of SARS-CoV-2 entry (*10, 11*). Other cellular proteases facilitate SARS-CoV-2 cell tropism, like furin, possibly at a post-binding step (*12, 13*).

Considering that SARS-CoV-2 may enter the cell by endocytosis, it would be partially acid pH dependent and should exit endosomes by fusion to start replication (*13*). SARS-CoV-2 replication occurs in the cytoplasm at double membrane vesicles of possible origin in the endoplasmic reticulum, which provide support to the viral RNA replication and transcription complex (*14-16*).

SARS-CoV-2 endosomal pH-dependent masking through endosomal cleavage was recently described (*8*). Similarly, endosomal processing of the Ebola virus (EBOV) glycoprotein trimer enhances infectivity and evidence a conformational masking of the receptor binding site of this protein (*17, 18*). A similar process has been described for the human immunodeficiency virus (HIV) envelope trimer (*19*). Therefore, the similarities found between these viruses were the start point of this work. EBOV enters the cell using the endocytic pathway, and its entry is mediated by the viral glycoprotein (GP), which decorates the viral surface organized in trimeric spikes (*20*). The GP is then processed by endosomal cathepsins to remove its heavily glycosylated C-terminal residues and the glycan cap. This removal produces the cleaved form of the N-terminal receptor binding subunit GP1 (GP_CL_) that is required to mediate fusion at the endosomal membrane. Two simultaneous publications described first that the endosomal receptor called NPC1 is an intracellular host receptor for EBOV (*21, 22*). Since then, the relevance of NPC1 on viral infections have been shown for different viruses including HIV (*23*), Hepatitis C (HCV) (*24, 25*), Chikungunya virus (CHIKV) (*26*) and several Flaviviruses such as Dengue (DENV) (*27, 28*) and ZIKA virus (*29*) among others (*26, 30*).

Given SARS-CoV-2 can be internalized via clathrin- and non-clathrin-mediated endocytosis; some researchers hypothesized a role for NPC1 in SARS-CoV-2 infection that remains to be determined (*31-34*). Due to the increasing relevance of this molecule, we designed our study to investigate the potential binding of SARS-CoV-2 proteins to NPC1, according to the existing evidence of SARS-CoV-2 endosomal passage (*35*).

## Materials and Methods

### Cell culture and viruses

Human embryonic kidney cells 293T/17 (HEK 293T; ATCC-CRL-11268) were cultured in Dulbecco modified Eagle medium (DMEM) at 37 °C and 5% CO_2_ atmosphere, supplemented with 100 IU/ml penicillin, 100 µg/ml streptomycin, 1X GlutaMAX (Thermo Fisher) and 10% heat-inactivated fetal bovine serum (FBS). Huh-7 Lunet C3 cells, a gift from T. Pietschman (Twincore, Germany), were cultured at 37 °C in Dulbecco’s modified Eagle’s medium (DMEM) supplemented with 100 IU/ml penicillin, 100 µg/ml streptomycin, 10mM HEPES, 1X NEAA and 10% of heat-inactivated fetal bovine serum (FBS).

For virus infections, we used common cold coronavirus 229E, which expresses the green fluorescent protein (GFP) gene (229E-GFP) (*36*). This recombinant virus was kindly given by V. Thiel, at the University of Bern, in Switzerland. The infection experiments were conducted at 33°C and 5% CO_2_.

### Design and construction of plasmid that express the SARS-CoV-2 N tag to EGFP

The methodology used in this part of the study was previously used in Garcia-Dorival et al., 2016 (*37*). To generate the SARS-CoV-2 N with N-terminal EGFP tag (EGFP-N), a codon optimized cDNA sequence for the ORF of SARS-CoV-2 N (NCBI reference sequence number: NC_045512) was cloned into the pEGFP-C1 (by GeneArt-Thermo Fisher Scientific). Once cloned, the sequence of the plasmid EGFP-N was confirmed by sequencing (Gene Art–Thermo Fisher Scientific).

### Expression of tagged-N protein and EGFP in HEK 293T cells

To transfect HEK 293T cells, four 60mm dishes were seeded with 2.5 ×10^6^ cells each 24 hours prior to transfection in DMEM complete medium described above. Then, a transfection of EGFP or EGFP-N was done using Lipofectamine 2000 (Thermo Fisher Scientific), following the instructions of the manufacturer. Twenty-four hours post transfection the cells were harvested, lysed and immunoprecipitated using a GFP-Trap kit (Chromotek).

### Immunoprecipitations (IP)

EGFP-N and EGFP immunoprecipitations (IP) were done using a GFP-Trap®_A (Chromotek). To do the IPs, the cell pellet was resuspended in 200μl of lysis buffer (10mM Tris/Cl pH 7.5; 150mM NaCl; 0.5mM EDTA; 0.5%NP40) and then incubated for 30 minutes on ice. The lysate was then clarified by centrifugation at 14000 *x g* and diluted five-fold with dilution buffer (10mM Tris/Cl pH 7.5; 150mM NaCl; 0.5mM EDTA). The GFP-Trap agarose beads were equilibrated with ice-cold dilution buffer and then incubated with diluted cell lysate overnight at 4°C on a rotator, followed by centrifugation at 2700 *x g* for 2 minutes. The bead pellet was wash two times with wash buffer (10mM Tris/Cl pH 7.5; 150mM NaCl; 0.5mM EDTA). After removal of wash buffer, the beads were resuspended in 100μl of Sample Buffer, Laemmli 2X Concentrate (Sigma Aldrich) and boiled at 95° for ten minutes to elute the bound proteins. Buffers used for Immunoprecipitations were all supplemented with Halt™Protease Inhibitor Cocktail EDTA-Free (Thermo Fisher Scientific).

### Co-Immunoprecipitation (Co-IP)

Similar to what was described in Garcia-Dorival et al, 2016 (*37*), Co-IP for NPC1 was performed using 50μl of the Immobilized Recombinant Protein G Resin (Generon) and specific antibodies against NPC1 (Abcam, ab108921). The cell pellets were incubated for 30 minutes on ice with 200μl lysis buffer. The lysate was then clarified by centrifugation and diluted five-fold with dilution buffer prior to adding 2μg of the primary antibody and then incubated at 4°C on a rotator for two hours. The protein G resin (Generon) were equilibrated with ice-cold dilution buffer and then incubated at 4°C on a rotator with diluted cell lysate containing the antibody overnight at 4°C on a rotator, followed by centrifugation at 2500 *x g* for 2 minutes to remove non-bounds fractions. The wash and elution steps were performed as describe previously in GFP co-immunoprecipitation.

### Western blot analysis

To do confirm the expression of GFP and GFP-N proteins, an SDS-PAGE and a western blot (WB) was done. For the SDS-PAGE, a Mini-PROTEAN TGX Gels were used (Bio-Rad 4561096), then, the gels were transferred to PVDF membranes using the Trans-Blot Turbo Transfect Pack (Bio-Rad 1704159) and the Trans-Blot Turbo system (Bio-Rad). Following this, the transferred membranes were then blocked in 10% skimmed milk powder dissolved in TBS-0.1% Tween (TBS-T) (50mM Tris-HCl (pH8.3), 150mM NaCl and 0.5% (v/v) Tween-20) buffer for one hour at room temperature. Primary antibody was diluted 1:1000 in blocking buffer and then incubated at 4°C overnight. After three washes, blots were incubated with appropriate HRP secondary antibody diluted in blocking buffer at a 1:5000 for 1 hour at room temperature. Blots then were developed using enhanced chemiluminescence reagent (Bio-Rad) and detected with ChemiDoc™ XRS Gel Imaging System using Image Lab™ software (Bio-Rad).

### Production of SARS-CoV-2 N protein in the baculovirus system

The sequence of the N protein published in the NCBI database was selected (GenBank accession number: 43740575 / NCBI reference sequence number: NC_045512). The codon usage of the N encoding gene was optimized for its expression in insect cells (OptimumGene™-Codon Optimization algorithm) and the coding sequence for this protein was synthesized by the company GenScript. The donor plasmid pFastBac1 containing an expression cassette expressing the recombinant protein under the control of the polyhedrin promoter was obtained. The Bacmid for the generation of the baculovirus was prepared in E. Coli DH10Bac bacterial cells containing the mini-Tn-7-replicon. Bacmids were transfected in the regulatory Sf9 cells and a viral clone selection was made by two rounds of plaque cloning to obtain the working virus stock. The baculovirus genome region was sequenced to determine the integrity of the N gene in the recombinant baculovirus named rBacN.

### SARS-CoV-2 N protein production in pupae

The production of SARS-CoV-2 N protein in insect pupae (*Tricoplusia ni; T. ni*) was performed as previously described (*38*). Briefly, pupae were allocated in the inoculation robot that dispensed a maximum of 5 μl with the baculovirus titers protein in 5 days pupae incubation time in constant temperature and humidity chambers. After that period, pupae were collected and stored frozen, before downstream processing. *T*.*ni* pupae containing the recombinant protein were homogenized in extraction buffer. Then, subsequent steps of clarification, diafiltration and His-tag purification were carried, out in order to obtain purified SARS-CoV-2 N protein. Protein concentration, yield and level of purity were determined by SDS-PAGE analysis using 4–20 % or 12 % Mini-Protean TGX precast gels from Bio-Rad. Gels were stained with QC Colloidal stain (3 ng sensitivity) in the case of concentration and yield evaluation and with SYPRO Ruby (1 ng sensitivity) in the case of level purity analysis, both from Bio-Rad. Recombinant SARS-CoV-2 N protein produced in pupae was measured by band densitometry with the ChemiDoc™ XRS Gel Imaging System using Image Lab™ software (Bio-Rad). A BSA standard curve was used for quantification.

### ELISA assays

High-binding 96-well ELISA plates (Nunc) were coated with 0.5 µg/well of purified SARS-CoV-2 N protein in carbonate/bicarbonate buffer 0.05 M pH 9.6 and allowed to bind over night at 4°C. Then, endogenous human NPC1 and HSP90 were purified using immobilized Recombinant Protein G Resin (Generon) and 4 µg of specific antibodies against NPC1 (Abcam, ab108921) or HSP90 (Enzo Life Sciences, ADI-SPA-835) respectively. All steps were performed as described in Co-IP assays. Serial dilutions of these endogenous NPC1 and HSP90 were added to the plate and capture was allowed to proceed for 1 hour at 37°C. After that, plates were washed with PBST (PBS 0.1%Tween20) and the binding of NPC1 to SARS-CoV-2 N protein was detected with a rabbit anti-NPC1 antibody (1:2000), revealed with an anti-rabbit-horseradish peroxidase (HRP) (1:2000) using a colorimetric substrate (OPD) and finally, quantified by absorbance at 492 nm in the EnSight multimode plate reader of PerkinElmer. Effect of the compounds on the binding was performed pre-incubating 50 µM and 100µM of each compound with NPC1 1h a 37°C before adding it to the plate.

### Compounds studied

All the compounds tested in this work have a purity ≥95% by HPLC. SC compounds were synthesized at Centro de Investigaciones Biológicas (CIB-CSIC) following described procedures. All these molecules were included in the MBC chemical library and some of them were previously characterized as potential inhibitors of the protein-protein interaction between NPC1 and EBOV-GP (*39, 40*). The compounds tested in this study are shown in Figure 2 and were resuspended in DMSO at 50 mM. Sulfides SC198 and SC073, and carbazole SC816, were used at working concentrations of 5, 50 and 50 µM; benzothiazepines SC397, SC593, SC567, at working concentrations of 75 µM and SC338 at 100 µM respectively. The first three compounds were shown previously to be active against EBOV while the others were inactive (*40*). Compounds MBX2254 and MBX2270 were used as gold standards as they have been reported to inhibit EBOV-GP/NPC1 interaction with high selectivity (*41*). These compounds were purchased from MolPort and used at concentrations of 75 µM and 25 µM, respectively. Class II cationic amphiphilic compound U18666A is a drug that blocks cholesterol flux out of lysosomes and also inhibits Ebola virus infections. It was acquired from Sigma-Aldrich and used at 10µM (*42*). Imipramine, a hydrophobic amine and FDA-approved antidepressant drug was acquired from Sigma-Aldrich and used at 25 µM (*26, 43*).

**Figure 1.**
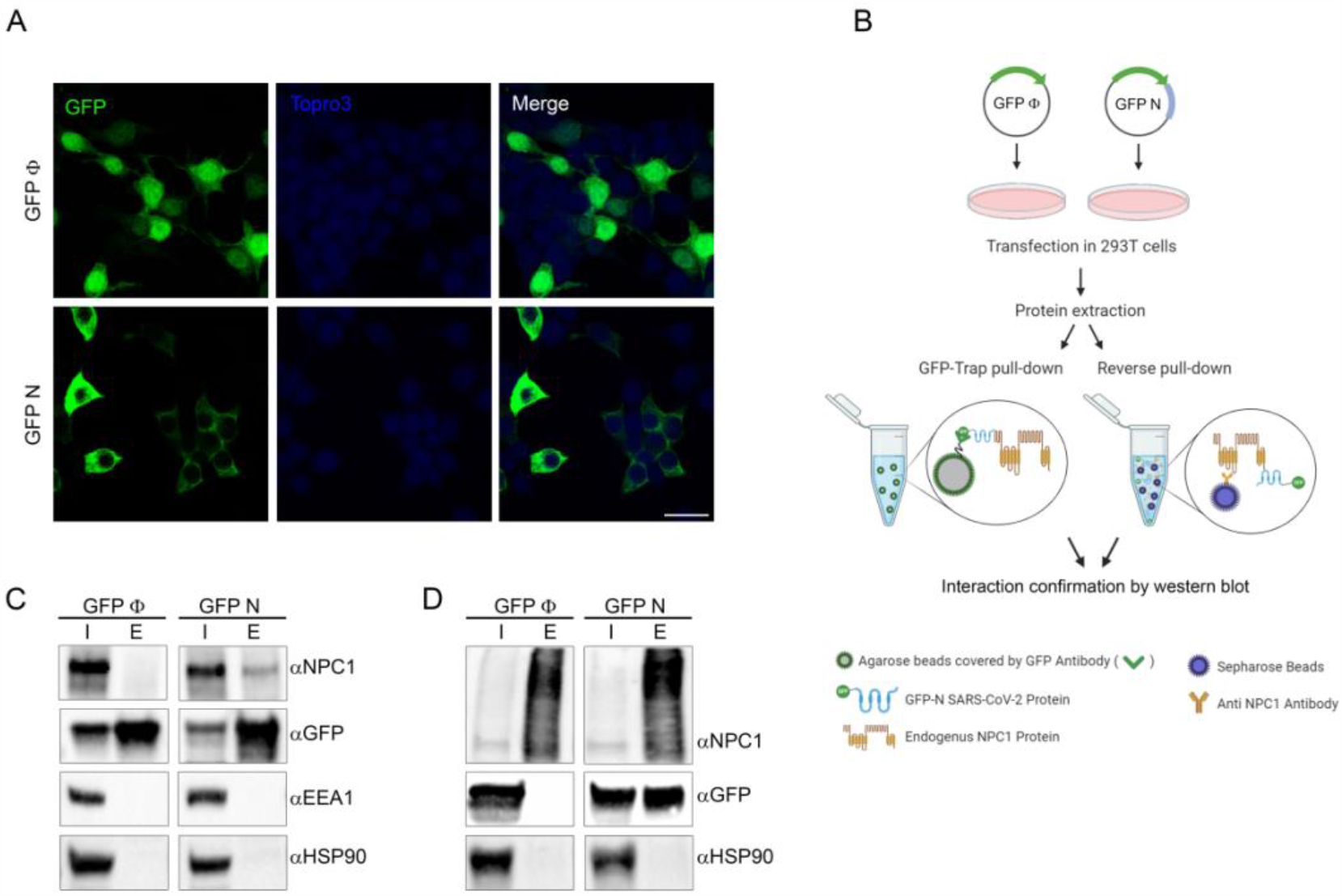
Immunoprecipitation analysis of SARS-CoV 2 N protein with endogenous NPC1. **A**. Expression of SARS-CoV-2 Nucleoprotein. Immunofluorescence of HEK 293T cells transiently expressing EGFP at the upper panel and EGFP-N protein at the lower panel showing different distribution as expected. GFP, Topro3 and Merge are indicated in upper panels in different colours. The scale bar indicates 25 µm. **B**. Schematic representation of the methodology used in this study for immunoprecipitation. **C, D**. Detection of SARS-CoV-2 N fussed to GFP, GFP control and cellular proteins analysed in the immunoprecipitation assay by western blot. **C**. Endogenous NPC1, EEA1, HSP90; and transfected EGFP-N and EGFP control were detected at the expected molecular weights. **D**. Endogenous NPC1, HSP90 and transfected EGFP-N and EGFP control were detected at the expected molecular weights from samples collected from the co-immunoprecipitations (reverse pulldown). Molecular weighs: NPC1∼175kD, EEA1∼110kD, HSP90∼75kD, EGFP-N∼70kD, EGFP∼27kD.

**Figure 2.**
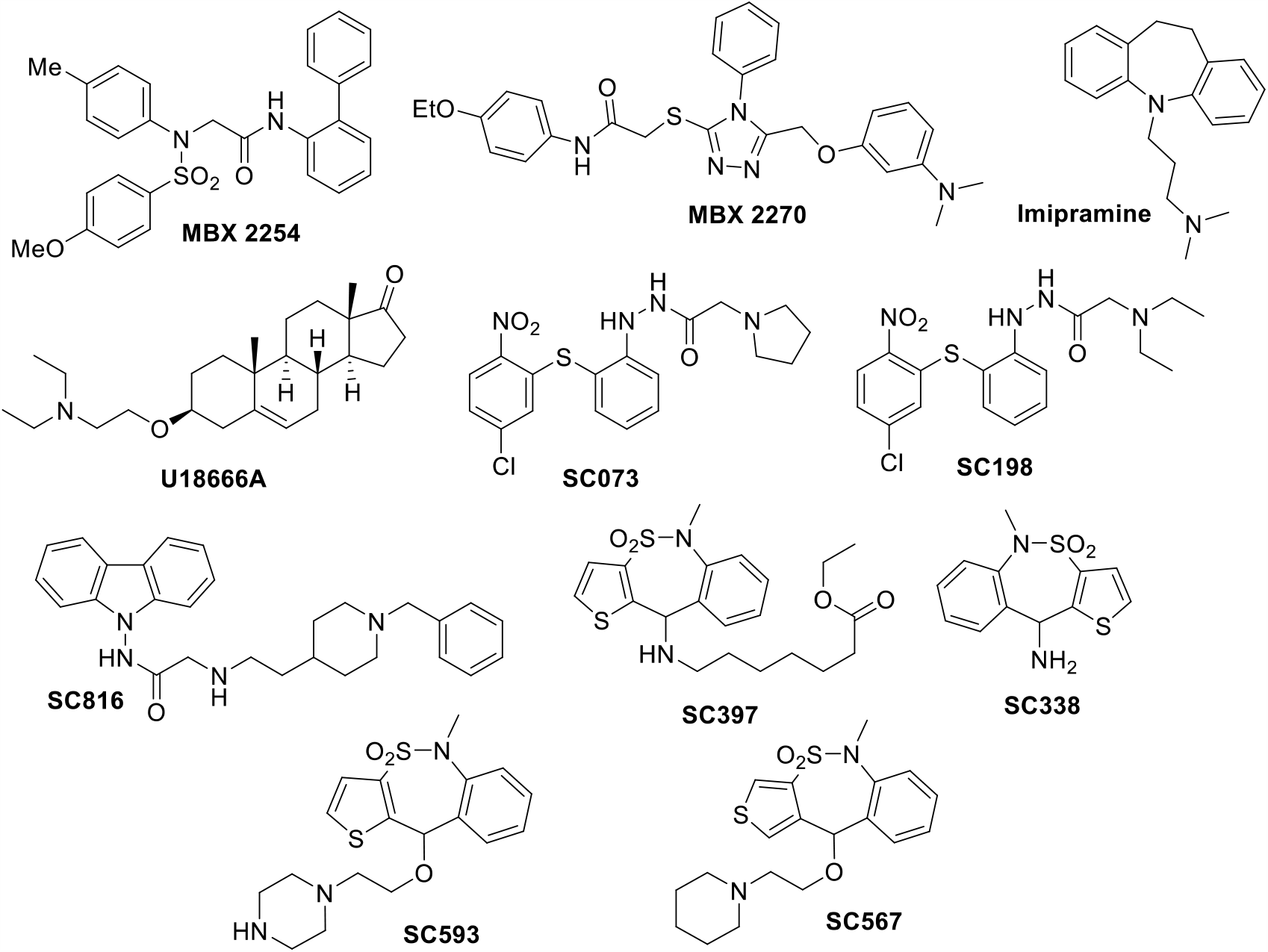
Chemical structure of small molecules used in this study.

### Cytotoxicity assays

Huh-7 cells were seeded in 96-well plates and incubated with DMEM containing each compound at concentrations ranging from 0 to 100 µM. After 24 hours, cell viability was measured by Cell Titer 96 AQueous Non-Radioactive Cell Proliferation Assay (Promega) following the manufacturer’s instructions. Absorbance was measured at 490 nm using an ELISA plate reader.

Cell viability was reported as the percentage of absorbance in treated cells relative to DMSO-treated cells (Figure S3). The 50% cytotoxic concentration (CC_50_) was calculated and non-toxic working concentrations (over 80% cell viability) used to test the activities of these compounds on CoV infection.

The values of the half maximal inhibitory concentration (IC_50_) inhibition of the infection presented on Figure 3 table correspond to the mean of 3 independent experiments. The IC_50_s values and dose-response curves were estimated using GraphPad Prism v6.0 with a 99% confidence interval.

**Figure 3.**
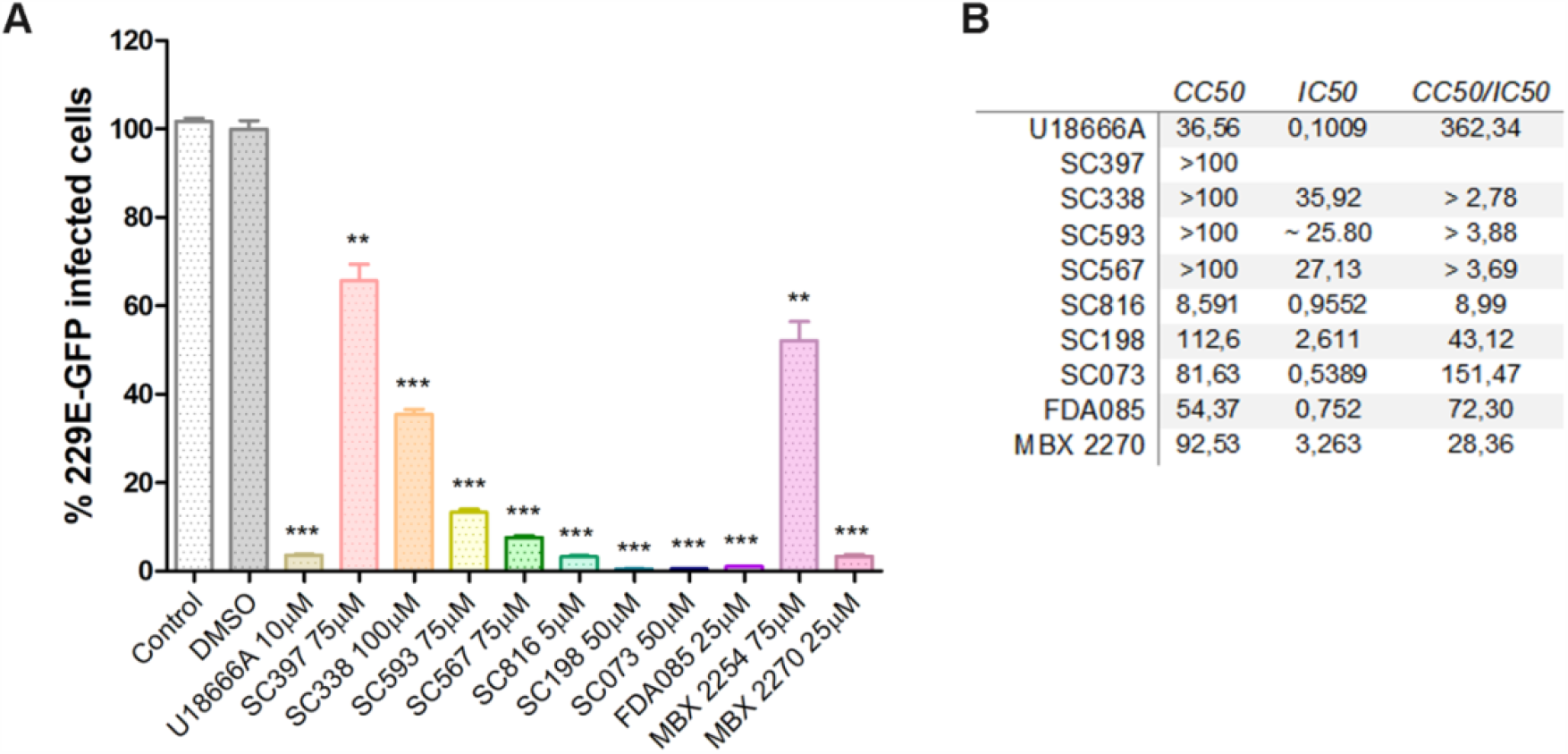
Activity of small-molecule inhibitors of NPC1 and related compounds against HCoV. **A**. Infectivity percentages of HCoV 229E-GFP in Huh-7 cells at 24 hpi. Y axis depicts GFP fluorescence intensity in controls and cells pretreated 1 h before infection with selected compounds at the concentrations above described (\*\**p* < 0.001; ***p < 0.0001). **B**. IC_50_ values (µM) were determined for these compounds. For determination of CC_50_ values, cells were treated with compound alone, and values (µM) were determined from linear portions of the dose-response curves shown in Supplementary Figure S4. SI, selectivity index (CC_50_/IC_50_).

### Flow cytometry analysis

Detection of CoV infected cells was performed by flow cytometry. Huh-7 cells were pre-treated with compounds at the indicated concentrations in growth medium for 1 h at 33 °C, followed by infection with 229E-GFP at a multiplicity of infection (MOI) of 1 pfu/cell for 24 h. Cells were washed twice with growth medium after 90 min of adsorption at 33°C, and incubated with DMEM 10% 24 h. Cells were then harvested with PBS-EDTA 5mM, and diluted in PBS. Detection of 229E-GFP infected cells was performed by analyzing GFP expression. In order to determine the percentage of infected cells per condition, 8,000 cells/time point were scored using FACS Canto II flow cytometer (BD Sciences) and analyzed using the FlowJo software. Untreated control infected cultures yielded 75–90% of infected cells from the total cells examined. Infected cell percentages obtained after drug treatments were normalized to DMSO values.

### Statistical analysis

The experimental data was analyzed by one-way ANOVA by Graph Pad Prism 6 software. For multiple comparisons, Bonferroni’s correction was applied. Values were expressed in graph bars as mean ± SD of at least three independent experiments unless otherwise noted. A *p* value <0.05 was considered as statistically significant.

## Results

### Interaction of SARS-CoV-2 N protein with NPC1

To investigate the interaction of SARS-CoV-2 Nucleoprotein (N) with NPC1, N protein was expressed as an EGFP-fusion protein in HEK 293T cells. Then, proteins were extracted from lysed cells and assayed for immunoprecipitation (IP) using a high affinity EGFP immunoprecipitation kit (GFP-Trap). Finally, protein expression/interaction was confirmed by fluorescence and western blot analysis (Figure 1A and 1C). This pipeline (Figure 1B) has been used to detect protein interacting partners for other RNA virus such as EBOV (*37, 44*). HEK 293T cells were selected due to their high efficiency of transfection, being the cell line of choice for protein-protein interaction studies of several viruses including SARS-CoV-2 (*45*).

Protein expression of EGFP-N and EGFP alone was confirmed using western blot analysis and fluorescence (Figure 1A and 1C). The efficiency of transfection for both plasmids was approximately 80% (Figure 1A). EGFP-N and EGFP were then immunoprecipitated using an EGFP-Trap. After immunoprecipitation, both input (cell lysate) and bound (or elution) samples were analysed by western blot. Proteins corresponding to the molecular weight of EGFP-N (70 kDa) and the EGFP empty (27 kDa) were detected using an anti-EGFP antibody (Figure 1C). NPC1, as an endogenous protein, was also detected in both input samples (EGFP-N and EGFP); but only in the bound fraction of EGFP-N sample (Figure 1C). This experiment was repeated three times to ensure reproducibility (Supplementary figure S1A).

To further validate a specific interaction between EGFP-N and NPC1, two cellular proteins were selected as negative controls. In this case, HSP90 chaperone and endosomal protein EEA1 were used as controls given the abundance of these proteins in cells (Figure 1C).

### Validation of SARS-CoV-2 N interaction with NPC1

Co-immunoprecipitations against NPC1 (or reverse pull down) were performed to confirm and further validate the interaction between SARS-CoV-2 N and NPC1. SARS-CoV-2 N was overexpressed in HEK 293T cells and then cellular extracts were analysed by co-immunoprecipitation using protein G-beads and specific monoclonal antibodies against NPC1 (Figure 1B). Bound samples obtained from the co-immunoprecipitations were then analysed by western blot, which confirmed the presence of SARS-CoV-2 N (Figure 1D). As a result of this interaction, we hypothesized that NPC1 might have an important function in virus biology.

### Functional assays

For an orthogonal characterization of the interaction, we used NPC1 inhibitor drugs to inhibit human coronavirus (HCoV) infection. To do this, Huh-7 cells were treated with the inhibitor compounds for an hour at different concentrations and then, infected with HCoV 229E-GFP recombinant virus at a MOI of 1 pfu/ml. Cells were then analysed at 16 hpi.

Inhibitors MBX2254 and MBX2270 (Figure 2) were selected as they target NPC1 with high selectivity and both have been described to inhibit HIV-pseudotyped-EBOV-GP binding to NPC1 (*41*). MBX2254 and MBX2270 were used at 75 and 25 µM, respectively. We also used imipramine, a Food and Drug Administration (FDA)-approved drug, that inhibits EBOV and other viruses due to its ability to induce a phenotype similar to NPC1 deficiency (*46*).

Finally, we assayed a set of compounds initially selected by virtual screening of the MBC chemical library in the EBOV-GP/NPC1 interaction (*40*). These compounds were previously found to inhibit infection with EBOV pseudotyped retrovirus and some of them - sulfides and carbazoles - were able to disturb the NPC1-GP interaction in an ELISA assay. Compounds were classified in three chemical classes, sulfides SC198 and SC073, and carbazole SC816 used at 5, 50 and 50 µM respectively; and benzothiazepines SC397, SC593, SC567 (Figure 2), that were used at 75 µM, or 100 µM of SC338. Noteworthy, sulfides and carbazoles were found to potentially act through inhibition of NPC1-GP interaction, while benzothiazepines do not affect this interaction (40). Based on these previous results, the three classes were included in this study for comparative purposes. As a reference, we used U18666A compound (10 µM), known to inhibit cholesterol transport function of NPC1 and the infectious entry of several viruses including EBOV and ASFV (*23, 26, 42*).

We detected that MBX2270 derivative potently inhibited HCoV infection (50% inhibitory concentration IC_50_= 3.26 µM, selectivity index 28.36; Figure 3). In general, our results yielded significant inhibition >99% of HCoV infection with the U18666A compound and imipramine treatment and with sulfides from the library compounds (Figure 3A). Others yielded over 80% of infectivity inhibition (except for SC397 and SC338; Figure 3B). IC_50_ was <1 µM in several sulfide compounds (SC073 IC_50_= 0.53 µM, Index 151.47), U18666A (IC_50_ 0.1 µM, Index 362.34) and imipramine (FDA085; IC_50_= 0.75 µM, Index 72.3). Full information of dose/response curves for these chemicals are included in Supplementary Fig. S4.

In addition to these functional experiments, we also tested the ability of these compounds to disrupt the NPC1/SARS-CoV-2 N protein interaction in an ELISA assay, as described in Materials and Methods. First, we tested increasing concentrations of NPC1 and control protein HSP90 in plates coated with SARS-CoV-2 N protein. We detected a positive reaction with increasing concentrations of NPC1, while negative control HSP90 remained unaltered (Figure 4A). Then, we analysed the inhibition of NPC1/SARS-CoV-2 N binding with a sample of the compounds previously described in this study. We obtained a significant inhibition of this specific binding in those samples tested with one inhibitor compound from each class 100 µM SC073 and 50 or 100 µM of MBX2270 (Figure 4B).

**Figure 4:**
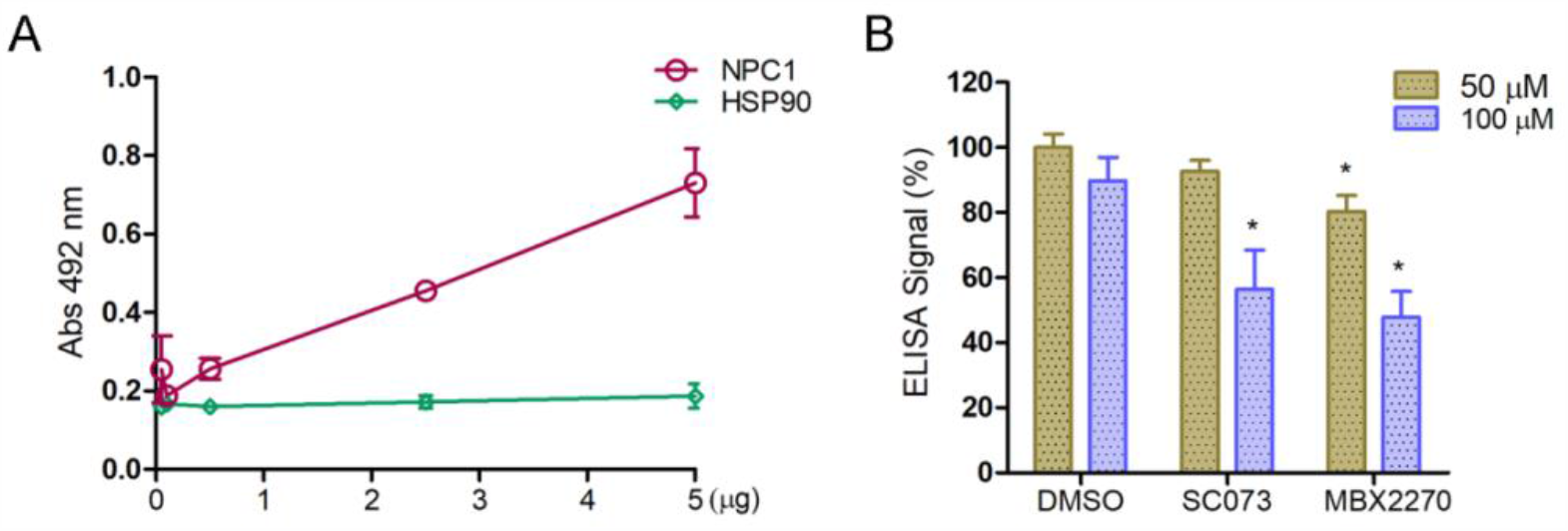
Inhibition of NPC1 binding to N protein by chemicals in an ELISA assay. **A**. Binding of increasing concentrations of NPC1 or HSP90 to N protein. HSP90 was used as a negative control. Concentrations analyzed were 5, 2.5, 0.5, 0.1 or 0.05 µg. **B**. DMSO or 50/100 µM chemical compounds were incubated with 5 µg of purified endogenous NPC1 before being added to ELISA plates previously coated with purified N protein (0.5 µg/well). Then, the binding of purified NPC1 protein to viral N protein was determined with an anti-NPC1 antibody revealed with an anti-rabbit-HRP. Absorbance was measured at 492 nm after addition of substrate. Percentages of binding were related to DMSO (\**p* < 0.01).

## Discussion

Current COVID-19 pandemic has affected millions of people all around the globe and has been one of the mayor challenges in this century due to a great loss of lives and significant economic losses (*1*). This highlights an urgent need for developing efficient therapeutics against SARS-CoV-2 since there is no licensed treatment available.

There are several pathogenic viruses, that are known to use the endocytic pathway to enter the cell, the most important being EBOV. Two publications described simultaneously that the endosomal protein called NPC1 or Niemann-Pick type C1 is a host receptor for EBOV (*21, 22*). EBOV entry is mediated by the viral glycoprotein (GP) which is organized in trimeric spikes at the viral surface (*20*). NPC1 binding requires the processing of viral GP. GP cleavage by endosomal cathepsins unmasks the binding site for NPC1 by removing heavily glycosylated C-terminal residues and the glycan cap to produce the cleaved form of the N-terminal receptor binding subunit GP1 (GP_CL_). Finally, GP_CL_-NPC1 binding within endosomes is required to mediate fusion and viral escape into the host cytoplasm (*21, 22*) as a second intracellular receptor (*47*).

Thereby, NPC1 could be used as an important druggable target (*22*) on viral infection. An example of this is compound U18666A, which blocks intracellular cholesterol efflux mediated by NPC1, along with imipramine severely impacts EBOV (*22, 42, 43*) and other viruses like HIV (*23*) and DENV (*27, 28*), CHIK (*26*), ZIKV (*29*) and other Flavivirus. Knockdown or chemical impairment of NPC1 severely reduced cholesterol supply at the Hepatitis C virus replication sites altering the replication membranous web (*25*).

SARS-CoV-2 infection starts with the interaction between spike glycoprotein (S) with the ACE2 cellular receptor. It requires activation by the TMPRSS2 at the plasma membrane. TMPRSS2 is located at the vicinity of ACE2 in lipid rafts and elicits plasma membrane fusion (*31*) that results severely impaired using chemical inhibitors of this protease (*10, 11*). Apart from TMPRSS2, the lysosomal proteases, specifically cathepsin L, are crucial for SARS-CoV-2 entry via endo-lysosomes (*11*). Both, TMPRSS2 and cathepsin L proteases have cumulative effects along with the cleavage caused by furin at the Golgi (subsequent to S-protein synthesis during viral packaging) on activating SARS-CoV-2 entry and penetration in the cytoplasm.

SARS-CoV-2 would traffic the endocytic pathway inside the early and late endosomal vesicles to finally fuse with lysosomes, an essential stage for viral uncoating and fusion (*31*). According to that, viral infection is abrogated by drugs interfering endosome acidification (*11*). Also, SARS-CoV-2 pseudovirions infection is inhibited using drugs targeting the late endosomal compartments, like cathepsin L, two-pore channel 2 (TPC2), or PIKfyve. Inhibitors against these proteins dramatically reduce infection, indicating that TPC2, cathepsin L, endosomal maturation, and endosomal acidic luminal pH, are crucial host factors for endocytosed SARS-CoV2 entry (*11, 35*). Thus, late endosomes/lysosomes are proposed as relevant organelles to develop therapeutic targets against infection by SARS-CoV-2 (*8, 11, 31-34, 48*).

A recent study discovered that SARS-CoV-2 non-structural protein 7 (nsp7) strongly interacts with Rab7a, and its depletion causes retention of ACE2 receptor inside late endosomes (*45*). Other reports highlighted the relevance of a variety of proteins involved in cholesterol biosynthesis, including NPC1 infection (*32*). Also, the cholesterol biosynthesis pathway is downregulated during SARS-CoV-2 infection and, according to that, drug treatments that regulate this pathway impact the infection (*49*).

Here, we described in this study an interaction between SARS-CoV-2 N protein and NPC1. This interaction unveiled a novel host-based target for antivirals and a potential host factor for SARS-CoV-2 infectivity. As in other viruses, this interaction could possibly regulate and modify cholesterol efflux from late endosomes and alter the lipid composition in cellular membranes in its own benefit (*50*). Besides, we presented data on how several compounds that block NPC1 function severely impact 229E HCoV infection in a functional assay, which suggests an essential role for NPC1 in HCoV infectivity.

Small molecule inhibitors were crucial to determine that NPC1 was essential for EBOV infection (*22*). Compounds MBX2254, an aminoacetamide sulfonamide, and MBX2270, a triazole thioether were reported to inhibit EBOV infection with high selectivity (*41*). All those compounds have NPC1 as a target and we found that those chemicals strongly inhibited HCoV 229E infection. Also, compounds found using the NPC1/EBOV-GP interaction for the screening of a library of compounds, namely sulfides, carbazoles and benzothiazepines (shown in Figure 2), were tested for HCoV inhibition. We have shown that compounds that inhibited EBOV-GP/NPC1 binding, namely sulfides SC073 and SC198 together with carbazole SC816, and not others, presented a potent inhibition of 229E-CoV infection. These chemicals have been shown to inhibit EBOV binding to NPC1 and the infection of EBOV-GP pseudovirions elsewhere (*40*). We have shown here that those compounds that were able to inhibit EBOV-GP/NPC1 binding were also capable to inhibit SARS-CoV-2 N protein/NPC1 binding in an ELISA assay.

To conclude, according to other authors and recent evidences (*31-34, 49*), we propose NPC1 as a potential therapeutic target for SARS-CoV-2 to combat COVID-19 pandemic. We show for first-time experimental evidences of the binding of SARS-CoV-2 Nucleoprotein (N) to NPC1. This important finding paves the way to direct medical efforts and therapeutics to NPC1 and to continue studies on cholesterol metabolism in SARS-CoV-2 infection.

## Acknowledgments

We are thankful to V. Thiel from the University of Bern, Switzerland for CoV 229E-GFP and T. Pietschman, Twincore, Germany for Huh-7 Lunet C3 cells. BioRender.com was used to created icons in Figures. This research was partially supported through “La Caixa” Banking Foundation (HR18-00469), Instituto de Salud Carlos III (ISCIII-COV20/01007), CSIC (201980E024 and 202020E079), Spanish Ministry of Science and Innovation (RTI2018-097305-R-I00) and the European Commission Horizon 2020 Framework Programme VACDIVA-SFS-12-2019-1-862874.

## Supplementary Figure Legends

**Supplementary Figure S1A:**
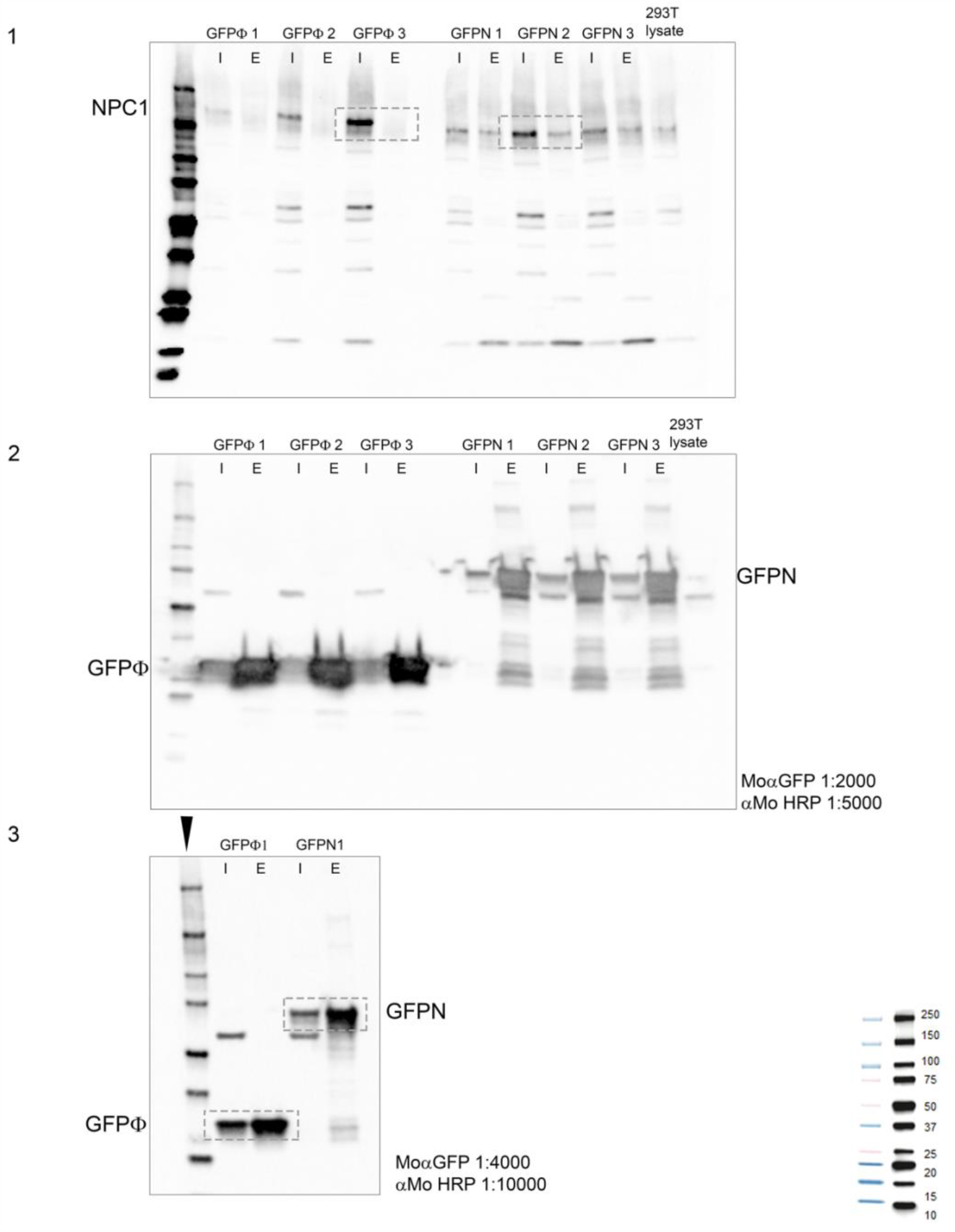
Membranes used to compose the figure 1C. Dashed boxes were taken to create the western blot composition showed in the figure. Membrane 1 triplicates revealed with rabbit anti-NPC1 antibody, membrane 2 triplicates revealed with mouse anti-GFP antibody and membrane 2 revealed with the same primary and secondary antibodies double diluted.

**Supplementary Figure S1B(cont):**
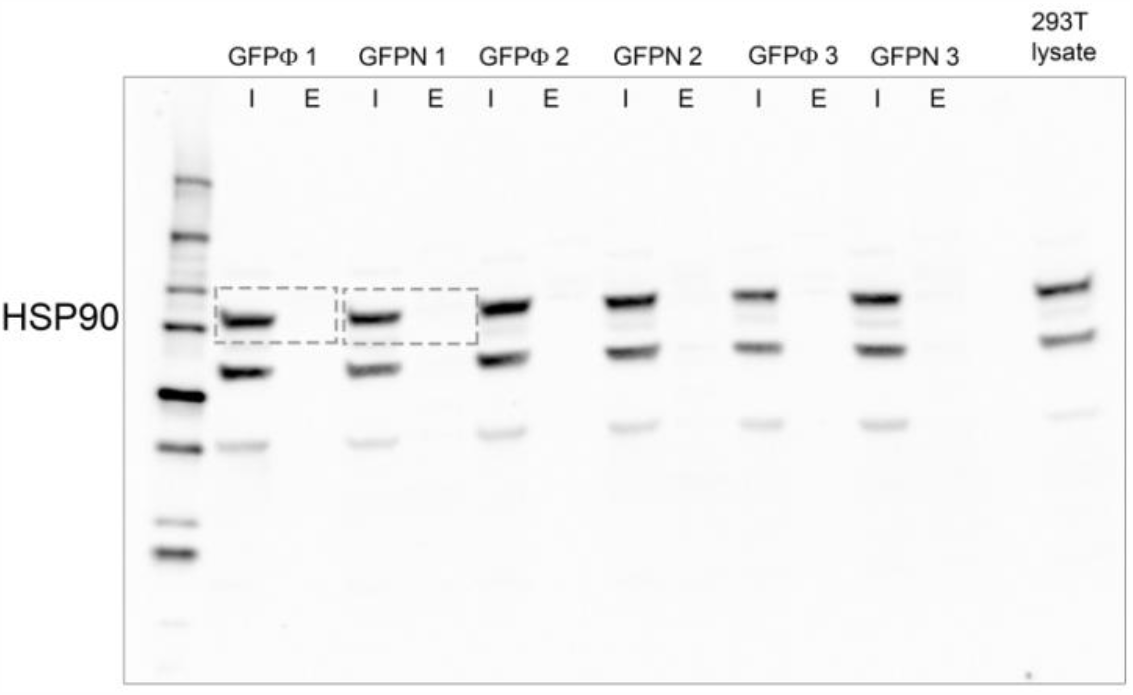
Membrane used to compose the figure 1C. Dashed boxes were taken to create the western blot composition showed in the figure. Original membrane.

**Supplementary Figure S2:**
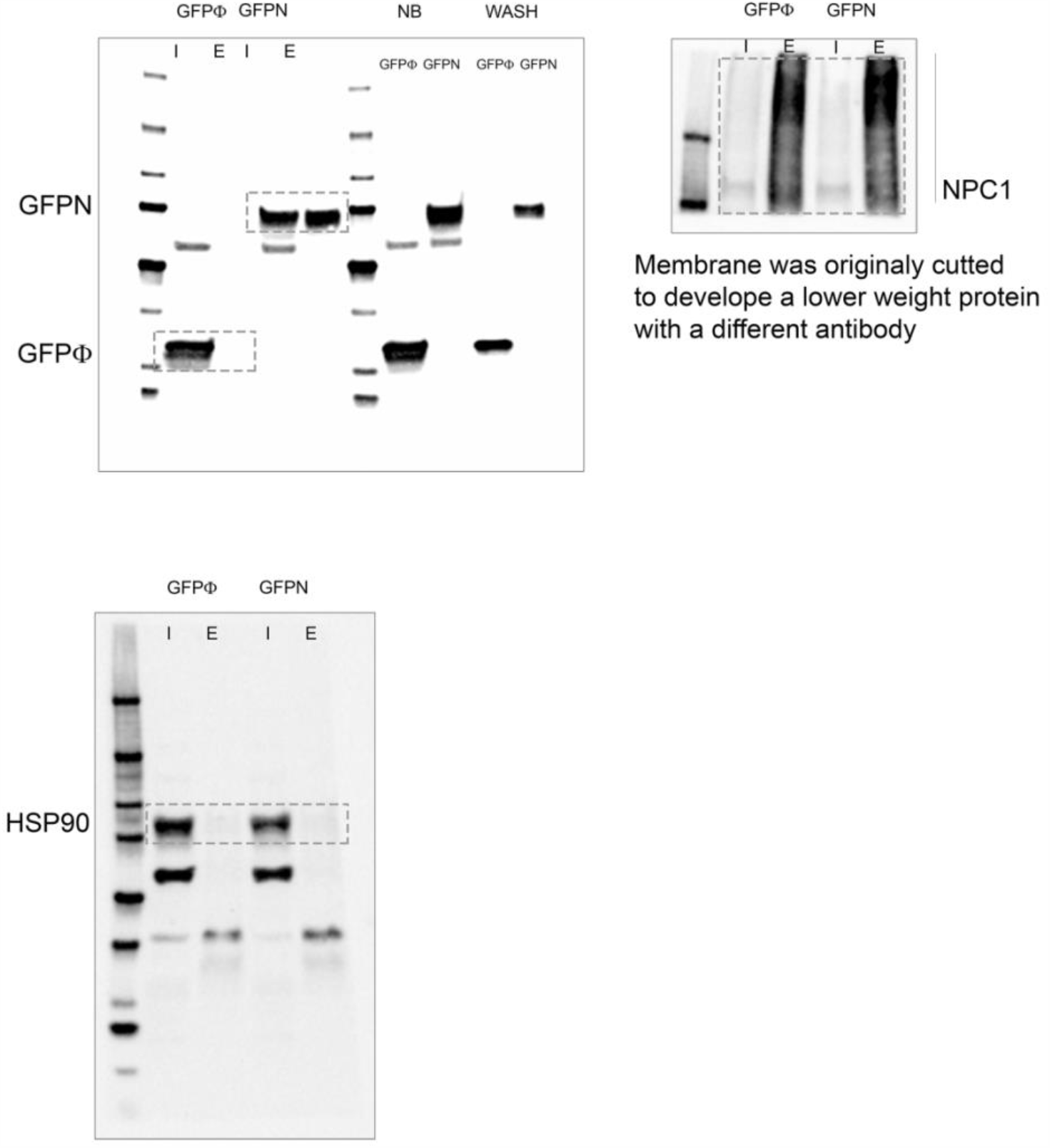
Membranes used to compose the figure 1D. Dashed boxes were taken to create the western blot composition showed in the figure. Original membrane 1 developed with Mouse anti-GFP antibody, membrane 2 developed with Rabbit anti-NPC1 antibody and membrane 3 developed with Rat anti-HSP90 antibody.

**Supplementary Figure S3:**
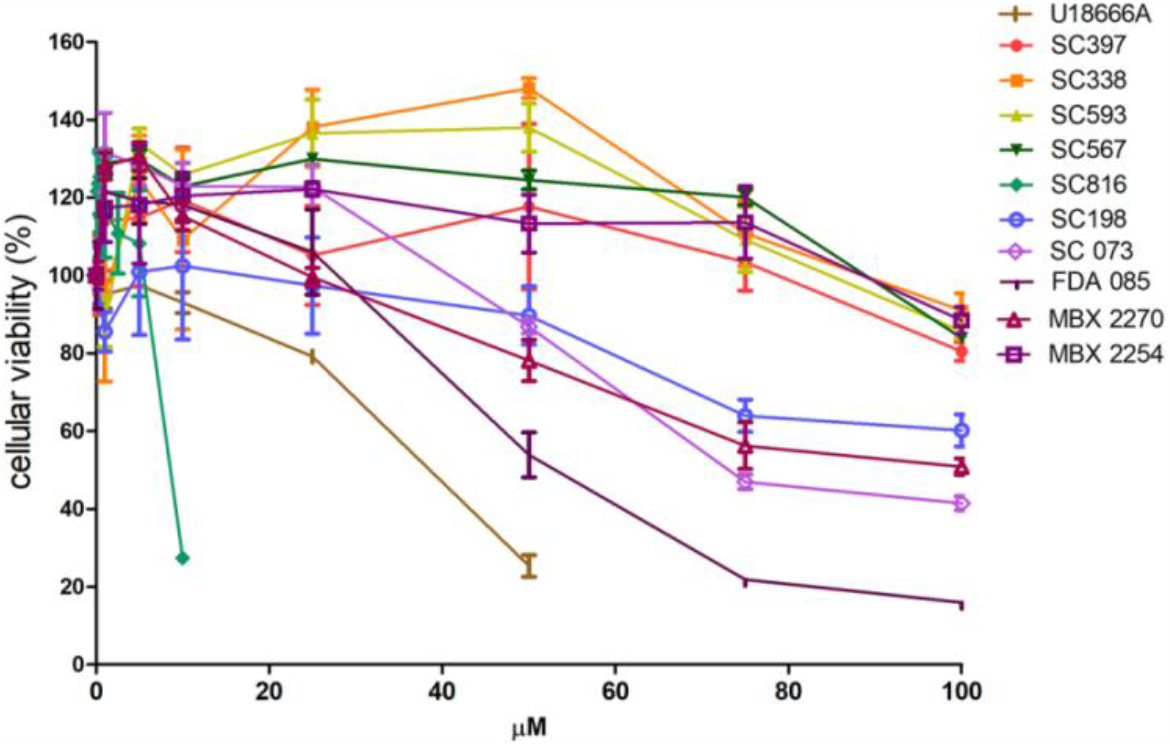
Cell viability under chemicals treatment. Cell viability measured after 24 hours incubation of Huh-7 cells with each compound at concentrations ranging from 0-100 µM in DMEM. Absorbance was measured at 490 nm using an ELISA plate reader. Y axis depicts median and standard deviations of the percentages of absorbance in compound treated cells relative to DMSO-treated cells.

**Supplementary Figure S4:**
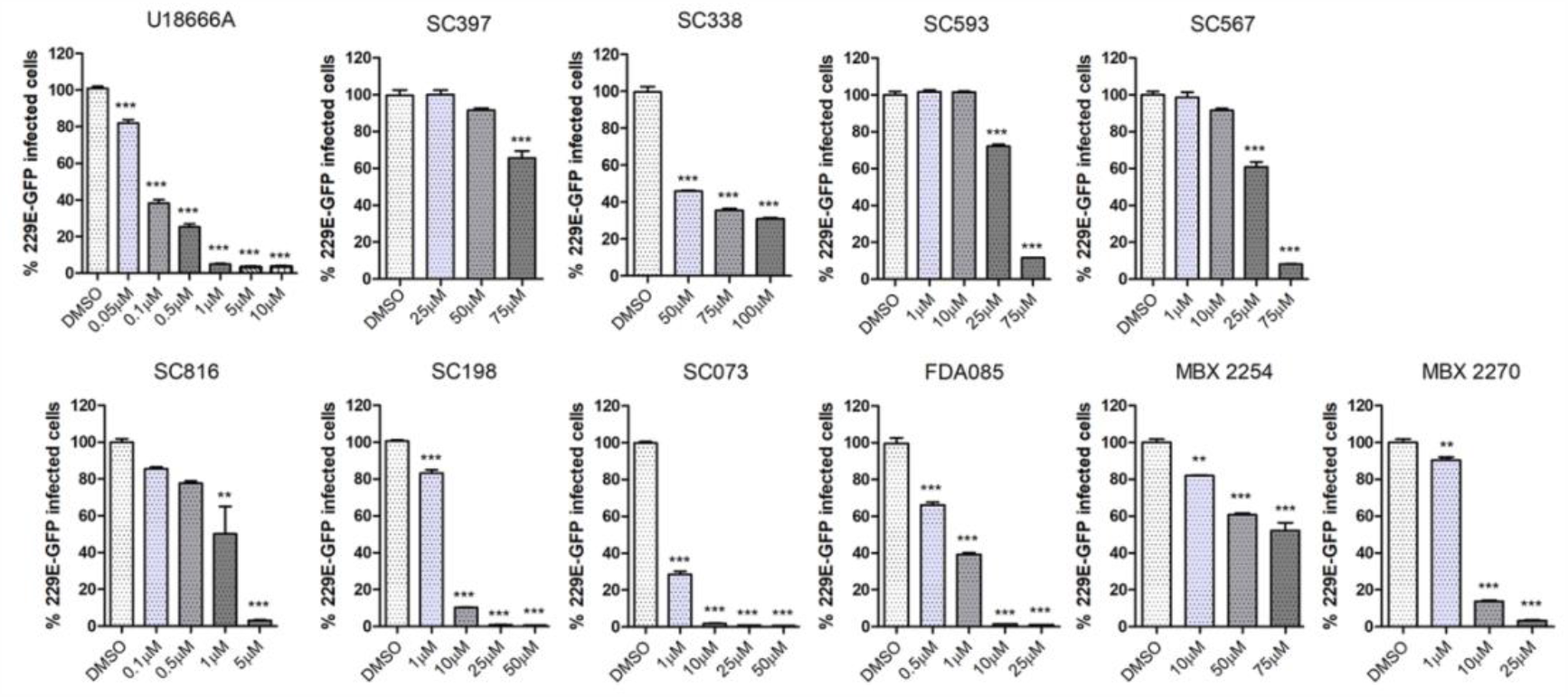
Dose-response curves of the compounds used. Dose-response curves with increasing concentrations of compounds at ranges selected depending on working concentrations for each compound.

## Notes

### Competing Interest Statement

The authors have declared no competing interest.

